# Meta-analysis of fecal metagenomes reveals global viral signatures and its diagnostic potential for colorectal cancer and adenoma

**DOI:** 10.1101/2022.07.17.500372

**Authors:** Fang Chen, Shenghui Li, Ruochun Guo, Fanghua Song, Yue Zhang, Xifan Wang, Xiaokui Huo, Qingbo Lv, Hayan Ullah, Guangyang Wang, Yufang Ma, Qiulong Yan, Xiaochi Ma

## Abstract

**Introduction:** Gut microbiome plays an important role in maintaining human health. Although mounting evidence has revealed the critical function of the gut bacteriome in the progression of CRC, the contribution of gut viral community to CRC is rarely studied.

**Objectives:** The present study aimed to reveal the gut virome signatures of colorectal adenoma patients and CRC patients and decipher the potential viral markers to build clinical predictive models for diagnosis.

**Methods:** 1,282 available fecal metagenomes data from 9 published CRC studies were collected. A new virus database was constructed based on a reference-independent virome approach for further analysis. Viral markers were filtered by statistical methods and used to build machine learning models such as Random Forest and Least Absolute Shrinkage and Selection Operator (LASSO) to distinguish patients from controls. New fecal samples were collected to validate the generalization of predictive model.

**Results:** The gut viral composition of CRC patients was drastically altered compared with healthy, as evidenced by changes in several Siphoviridae viruses and a reduction of Microviridae, whereas the virome variation in adenoma patients was relatively low. The viral markers contained the phages of *Porphyromonas*, *Fusobacterium*, *Hungatella*, and *Ruminococcaceae*. In 9 cohorts and independent validation cohorts, a random forest (RF) classifier and LASSO model got the optimal AUC 0.830 and 0.906, respectively. While the gut virome analysis of adenoma patients identified 88 differential viruses and achieved an optimal AUC of 0.772 for discriminating patients from controls.

**Conclusion:** Our findings demonstrate the distinctly different composition of gut virome between healthy controls and CRC patients, and highlight the potential of viral markers for clinical diagnosis.

## Introduction

Colorectal cancer (CRC) is one of the most common cancers with an increasing global incidence and mortality rate, especially in many low- and middle-income countries(*1*). Numerous studies have implicated gut bacteria in the development of CRC(*2–4*). According to next-generation sequencing, *Fusobacterium nucleatum* was enriched in the human colon cancer tissues and stool samples from the CRC patients compared to the healthy controls(*5*). *F. nucleatum* recruited tumor-infiltrating myeloid cells in the mouse model, which accelerated carcinogenesis(*5*). A recent study found that *F. nucleatum* contributed to intestinal tumorigenesis via promoting CRC cell glucose metabolism(*6*). Thomas *et al*. identified 25 accurate CRC biomarker species including *Parvimonas micra*, *Gemella morbillorum*, *Peptostreptococcus stomatis*, *Solobacterium moorei* and *Porphyromonas asaccharolytica* in human fecal metagenomes, through meta-analysis of cross-cohort studies(*2*). In certain studies, a prediction model based on these biomarkers demonstrated good discrimination(*2, 3, 7*). These results revealed multiple CRC-associated bacteria and provided insight into their potential pathogenic mechanisms in CRC.

Viruses are crucial members of the microbial ecosystem in the human gut and are often underemphasized in early studies. In recent years, many studies have determined that several human systemic diseases, such as rheumatoid arthritis and inflammatory bowel disease(*8–11*), were associated with both gut bacteria and viruses. Although some studies pointed out the association between specific viruses and CRC(*12–15*), only a few of them involved in the gut virome of CRC so far. Hannigan *et al*. reported that alpha and beta diversities of gut virome were not significantly different between CRC patients and healthy controls(*16*). The relative abundance of multiple gut viruses had a significant difference between CRC patients and health controls(*16–18*). In addition, prediction models based on CRC-associated viruses presented an acceptable potential for classifiability of controls vs. patients (∼0.802) in three studies(*16–18*). Among three studies, the viral markers in Nakatsu *et al.*’s study failed to correctly classify CRC patients vs. health controls among additional three independent cohorts(*17*). The reproducibility and predictive accuracy of microbial markers cannot be validated across multiple studies. This is because many biological confounders (e.g., host clinical parameters) can lead to false positives and the heterogeneity of data generation and processing can decrease the stability and reliability of results. To reduce the effect of biological and technical factors, previous studies performed meta-analyses on the gut bacteriome and identified CRC-associated changes that were consistent across populations(*2, 3*). However, there is still lack of large-scale and cross-cohort studies about the CRC gut virome. Thus, it is crucial to perform meta-analyses across studies to avoid biased associations between the gut virome and CRC.

In this study, we downloaded 1,282 fecal metagenomes from 9 published cohorts, generated a nonredundant viral catalog, and profiled the gut viromes of each to characterize the gut viral signatures in CRC. By referring to the method of the previous studies(*2, 3*), we performed meta-analyses to identify accurate CRC-associated viral biomarkers across 9 cohorts of CRC patients and healthy controls. To figure out the diagnostic potential of these viral biomarkers for CRC, we performed intra-dataset and cross-dataset prediction and validation based on 9 cohorts. We further validated the efficiency of viral biomarkers in a newly recruited cohort and an additional independent cohort published recently.

## Material and methods

### Data collection

The publicly available datasets of 9 CRC studies, covering fecal metagenomes from 554 CRC patients, 182 adenomas patients, and 546 healthy controls, were downloaded from the NCBI SRA database using the following accession IDs: PRJEB6070 for Zeller_2014, PRJEB10878 for Yu_2015, PRJEB7774 for Feng_2015, PRJEB12449 for Vogtmann_2016, PRJNA389927 for Hannigan_2018, PRJNA447983 for Thomas_a_2019 and Thomas_b_2019, PRJEB27928 for Wirbel_2019, and PRJDB4176 for Yachida_2019. The metadata of samples were obtained from the basic research studies or extracted from the NCBI BioSample database. Fecal metagenomes of the recently published validation cohort, Yang_2021, were downloaded from the NCBI SRA database with accession ID PRJNA763023.

### Recruitment of an independent cohort

To verify the accuracy of our virus marker model in other data, a newly recruited independent cohort was adopted as the validation cohort. Individuals were recruited at Dalian University Affiliated Xinhua Hospital and Dalian Medical University between 2020 and 2021. A total of 27 CRC patients and 28 healthy controls were included.

### Ethics statement

Written informed consents were obtained from all participants in this study. Ethical approval for this study was obtained from the Ethics Committees in the Dalian University Affiliated Xinhua Hospital [Approval no. 2022-04-01]. All procedures followed were in accordance with the Helsinki Declaration of 1975, as revised in 2008.

### Metagenomic sequencing

DNA was extracted from fecal samples using a TIANamp Stool DNA Kit (TIANGEN, China). The quality of DNA was assessed with Qubit 2.0. The extracted DNA samples were stored at -80□ until use. A sequencing library was generated using the NEB Next® Ultra™ DNA Library Prep Kit (NEB, USA) following the manufacturer’s recommendations and index codes were added to each sample. Library quality was confirmed with an Agilent 2100. The clustering of the index-coded samples was performed on a cBot Cluster Generation System using the Illumina PE Cluster Kit (Illumina, USA) according to the manufacturer’s instructions. After cluster generation, the DNA libraries were sequenced on the Illumina NovaSeq platform and 150 bp paired-end reads were generated.

### Preprocessing and assembly

Raw reads were qualified via fastp v0.20.1 ^(*35*)^ with the options ‘-u 30 -q 20 -l 90 -y --trim_poly_g’, and human reads were further removed by matching quality-filtered reads against the human genome GRCh38 with bowtie2 v2.4.1 ^(*36*)^. The remaining clean reads of each sample were assembled into contigs using Megahit v1.2.9 with the options ‘--k-list 21,41,61,81,101,121,141’(*37*).

### Analyses of viral sequences

All assembled contigs (≥5 kb) were used to identify viral sequences in each sample. The detection, decontamination, and clustering of viral sequences were performed as described in our previous study(*38*). Taxonomic and functional annotation of viral sequences were also implemented based on the criteria described in our previous study(*38*). Virus-host prediction was performed based on the Unified Human Gastrointestinal Genome (UHGG) database(*27*) using two methods that included CRISPR-spacer matches and prophage blasts (see the methods of our previous study for the detailed flow(*38*)).

### Taxonomic profiling and diversity

Clean reads in each sample were mapped to the non-redundant viral catalog of 37,030 vOTUs using bowtie2 with the options ‘--end-to-end --fast --no-head --no-unal --no-sq. The abundance profile of vOTUs in each sample was generated by aggregating the number of reads mapped to each vOTU. Random subsampling of each sample was implemented to achieve a parity number of reads that equals the minimum number of mapped reads among all samples. After subsampling, the relative abundance of vOTUs was its abundance divided by the number of total mapped reads in each sample. The relative abundance profile at the family level was generated by aggregating the relative abundance of vOTUs assigned to the same family. Alpha diversity indexes were assessed based on the relative abundance profiles at the vOTU level. Shannon index was calculated using the function *diversity* with option ‘index = shannon’ in the R platform. The number of observed vOTUs was the count of unique vOTUs in each sample. In addition, the taxonomic profiling of the bacteriome was performed based on 4,644 prokaryotic genomes from UHGG(*27*) by the aforementioned methods. Alpha diversity indexes of the bacteriome were calculated based on the relative abundance profiles at the species level.

### Identification of the biomarkers

Based on vOTUs profile, the statistically different vOTUs among the groups were selected from each of nine public fecal shotgun CRC datasets by a Wilcoxon rank-sum test. Among these, the vOTUs existed in more than four datasets with consistent trend between patients and controls were screened. To overcome the limitations of single research, features were further filtered by meta-analysis. We converted vOTUs relative abundances to arcsine-square root-transformed proportions and used the *escalc* function from the R *metafor* package that employs Hedges’ g standardized mean difference statistic to calculate pooled effect size by random effects model. We assessed between study heterogeneity by using Cochran’s Q test and I^2^ index. P values obtained from the random effects models were corrected for multiple hypothesis testing using the Benjamini–Hochberg procedure, with FDR < 0.01 considered statistically significant. Within each cohort, we fitted a linear model with the R *limma* package for each vOTUs feature “log10(vOTUs feature) ∼ disease_state + age + BMI + gender”. Disease state corresponds to whether an individual was a case (2) or a control (1). The relative abundances of the vOTUs were logged as we found their distributions to be primarily log-normal. After getting the coefficient and corresponding standard error, the meta-analysis was performed to calculate the pooled coefficient and their 95% confidence interval. To avoid confounding due to intra-individual variation, potential confounders such as age, sex, and BMI were included in the models. Only the pooled coefficients with 95% CI that did not contain zero after adjusting for confounders would be kept as final CRC biomarkers.

### Correlation network analysis

We used three methods to evaluate whether there is a relationship between CRC-associated vOTUs and prokaryotes. 1) The aforementioned host assignment. 2) Based on the SparCC algorithm(*39*), co-abundance relationships were established on the read count profiles of vOTUs and prokaryotes using *fastspar* v0.0.10 (*40*) with the option ‘--iterations 20’, while *fastspar_pvalues* v0.0.10 was used to calculate p-value by 1,000 bootstraps datasets derived from *fastspar_bootstrap* v0.0.10. Co-abundance relationships with the threshold of the correlation coefficient > 0.35 and q-value < 0.01 (p-value adjusted by the function *p.adjust* with the option “method=BH”) were retained. 3) To identify co-occurrence relationships, the presence and absence of each vOTU and prokaryote were treated as binary traits in each sample, and then compiled into a contingency table that displayed the numbers of microbial populations. The pairwise co-occurrence relationship was assessed based on the contingency table using Fisher’s exact test by the function *fisher.test* in R platform, while p-value was adjusted by the function *p.adjust* with the option “method=BH” in R platform. If the odds ratio of co-occurrence pair was greater than 100 and the q-value was less than 0.0001, the two microbial populations were considered to be a co-occurrence. Finally, these relationships between CRC-associated vOTUs and prokaryotes were visualized using Cytoscape v3.8.2(*41*).

### Prediction modeling

Random forest model (*randomForest* package in the R platform, ntree=2,000) and L1-regularized (LASSO) logistic regression model (SIAMCAT(*42*) in R platform, the same parameter setting derived from Thomas’s study(*2*)) were used to build the classifier based on the vOTUs abundance profile. In LASSO, we performed log10-transformed (after adding a pseudocount of 1 × 10^-8^ to avoid non-finite values resulting from log10(0)) in the profile, and finally standardized it as z-scores. In random forest model, the profile was used without any pretreatment process. Data were split into training and testing sets for 5 times repeated, fivefold stratified cross-validation (each fold contained a balanced proportion of positive and negative cases). For each split, a machine learning model was trained on the training set, which was then used to predict the test set. Models were then evaluated by calculating the AUROC based on the posterior probability for the CRC class. In inter-cohort prediction, models built from one dataset are used to predict the other eight datasets. In the LOCO setting, one cohort serves as the test set and the remaining eight datasets served as the training set by using five times-repeated fivefold stratified cross-validations.

### Data and code availability

The raw metagenomic sequencing data of the independent cohorts have been deposited to the China Nucleotide Sequence Archive under the accession codes CNP0002641. The authors declare that all other data supporting the findings of the study are available in the paper and supplementary materials, or from the corresponding authors upon request.

## Results

### Collection of datasets

In this meta-analysis, we included 9 published datasets that used whole-metagenome shotgun sequencing to characterize the fecal microbial communities of patients with colorectal cancer or adenoma (Table 1; Table S1). Participants from all studies were diagnosed by colonoscopy or alternative methods, and the controls were confirmed after the absence of disease. Healthy subjects with a history of colorectal surgery in Yachida *et al.*’s study were excluded. Metagenomic data of these studies were processed using a uniform protocol to ensure consistency, and fecal metagenomes were excluded if the proportion of human DNA sequences exceeded 10% or the number of high-quality reads was less than 4 million. A total of 1,282 samples, including 554 colorectal carcinoma (CRC) patients, 182 adenoma patients, and 546 healthy controls, containing nearly 5.9 Tbp of data, were retained for further analysis (Table 1).

**Table 1.** Summary of sample characteristics of data sets included in this study.

| Dataset | Country | Number of samples |  |  | Data | Total | NCBI<br>accession ID |
| --- | --- | --- | --- | --- | --- | --- | --- |
|  |  | CRC | Adenoma | Control | per | data |  |
|  |  |  |  |  | sample<br>(Gbp) | amount<br>(Gbp) |  |
| Zeller_2014(19) | France | 33 | 33 | 35 | 1.5±1.2 | 152.8 | PRJEB6070 |
| Yu_2015(20) | China | 74 | - | 54 | 4.6±1.0 | 581.9 | PRJEB10878 |
| Feng_2015(21) | Austria | 46 | 47 | 63 | 4.2±0.7 | 654.4 | PRJEB7774 |
| Vogtmann_2016(22) | USA | 23 | - | 16 | 1.1±0.3 | 43.2 | PRJEB12449 |
| Hannigan_2018(16) | USA,<br>Canada | 16 | 8 | 16 | 0.8±0.3 | 30.0 | PRJNA389927 |
| Thomas_a_2019(2) | Italy | 29 | 27 | 24 | 4.6±2.7 | 365.5 | PRJNA447983 |
| Thomas_b_2019(2) | Italy | 32 | - | 28 | 3.9±1.7 | 235.0 | PRJNA447983 |
| Wirbel_2019(3) | Germany | 43 | - | 60 | 2.5±1.3 | 259.5 | PRJEB27928 |
| Yachida_2019(4) | Japan | 258 | 67 | 250 | 6.3±1.6 | 3,535.3 | PRJDB4176 |
| **Overall** |  | **554** | **182** | **546** | **4.6±2.3** | **5,857.5** |  |

### Diversity and overall structure of gut viral community in relation to CRC and adenoma

To characterize the gut viral communities, we performed *de novo* assembly of the fecal metagenomes (generating totaling 5.5 million contigs at a minimum length of 5,000 bp; Table S1) and identified 208,048 viruses from the assembled contigs based on the homolog-based and machine learning-based methods (see Methods). A nonredundant viral catalog of 37,030 viral operational taxonomic units (vOTUs; average length 30,111 bp, N50 length 44,627 ranging from 5,000 bp to 410,947 bp; Fig. S1A) were then generated under the species-level nucleotide similarity threshold of 95%(*23, 24*). The quality levels of these vOTU sequences were estimated by CheckV(*25*), which resulted 6.7% complete, 21.1% high-quality, 19.3% medium-quality, and 52.8% low-quality viral genomes, and 0.2% quality-undetermined sequences (Fig. S1B). Taxonomically, 45.9% of these vOTUs were assigned to a known viral family, the majority of them consisted of *Siphoviridae*, *Myoviridae*, *Podoviridae*, *Quimbyviridae*, and *crAss-like* viruses (Fig. S1C). The currently available collections of human gut virome, including the Gut Virome Database(*23*) (covering 13.97% vOTUs in this study), Gut Phage Database(*24*) (covering 35.72% vOTUs in this study), and Metagenomic Gut Virus catalog(*26*) (covering 25.93% vOTUs), identified 43.86 percent of 37,030 vOTUs (Fig. S1D).

Two parameters, the observed number of vOTUs and the Shannon index were used to estimate the within-sample (alpha) diversity of gut viral community of 9 analyzed datasets. The observed number of vOTUs in the virome of CRC patients was significantly higher in two datasets (Wilcoxon rank-sum test, p<0.001 in Feng_2015, *p*=0.006 in Thomas_b_2019), but showed no consistent trend in other studies; while the Shannon index showed no significant difference between two groups in all datasets. (Fig. S2A). Likewise, both diversity parameters were approximately equal in the viromes of adenoma patients and healthy controls (Fig. S2B). Furthermore, we found that the viral diversity parameters of samples were highly consistent with their bacterial diversities (Fig. S2C-D), suggesting an extensive connection between the virome and the bacterial microbiome.

Principal coordinates analysis (PCoA) showed a substantial difference in viral composition among the nine study populations (permutational multivariate analysis of variance [PERMANOVA] R^2^=11.9%, *p*<0.001; Fig. 1A). Despite that, the disease status of subjects still had a significant impact on the overall viral composition (PERMANOVA R^2^=0.8%, *p*<0.001). We then quantified the effect size of disease status on gut virome within each study and discovered that, with the exception of Hannigan et al.’s study, CRC status was significantly associated with viral composition in almost all cohorts (Fig. 1B). However, the adenoma status had no significant effect on viral composition in all cohorts (Fig. 1C). Furthermore, because individual heterogeneity has been shown to closely correlate with viral taxon variants, we assessed the effect sizes of host characteristics such as age, gender, and body mass index (BMI) on gut virome based on all data. BMI showed a considerable impact on the overall viral composition (PERMANOVA R^2^=1.1%, *p*<0.001), whereas the effect sizes of age and gender were relatively lower (R^2^=0.4% and 0.3%, respectively; Table S2). Moreover, adjusting the host’s age, gender, and BMI didn’t visibly change the size of the effect of disease status on the gut virome, suggesting that there was little interaction between them.

**Fig. 1.**
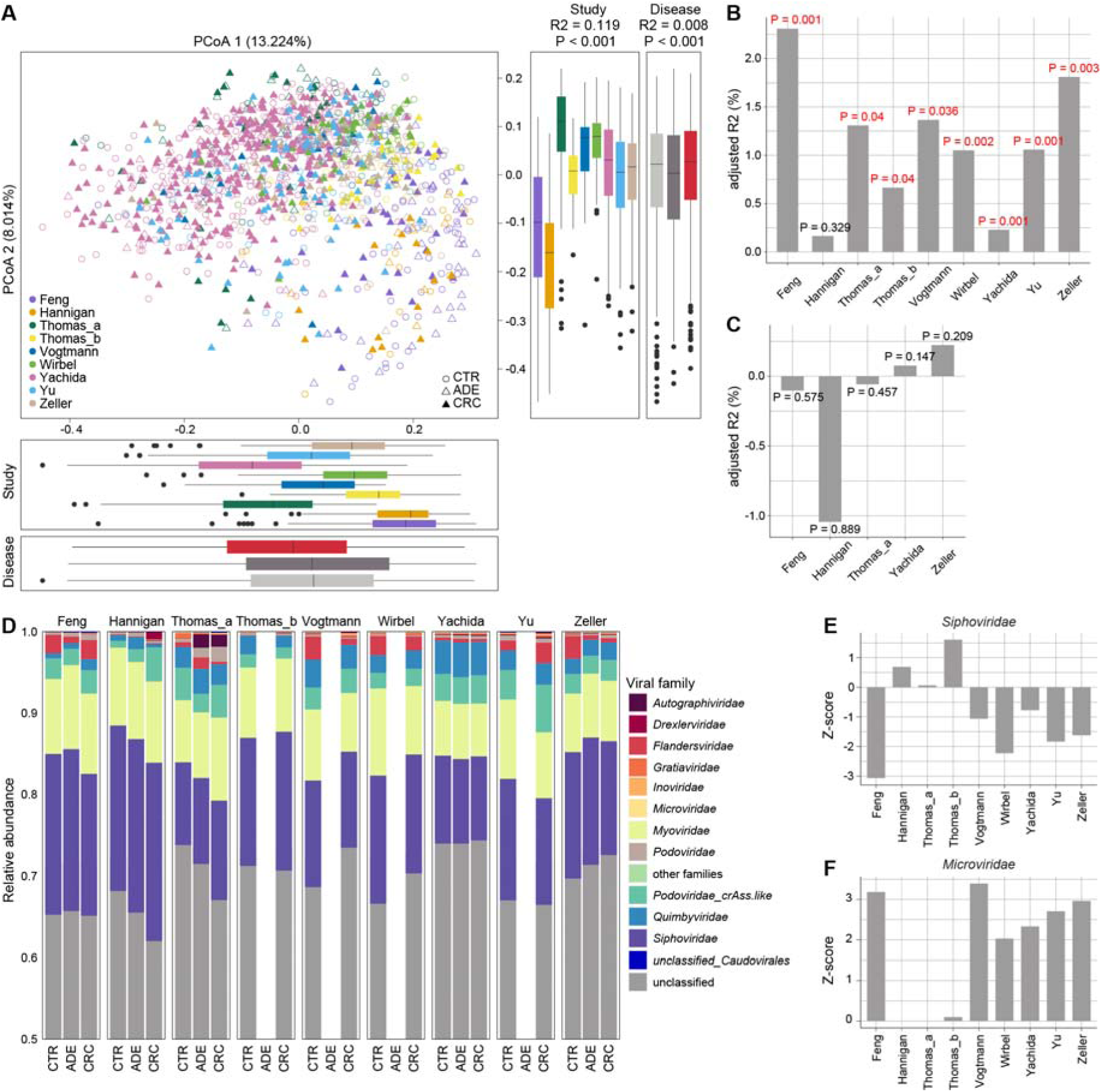
Viral community variation among the nine study populations. **(A**) Principal coordinates analysis (PCoA) based on the Bray-Curtis distance at the vOTU level. CTR, healthy controls; ADE, adenoma patients; CRC, colorectal carcinoma patients. Effect size (R2) and statistical significance were obtained by PERMANOVA (adonis). (**B** to **C)**, Effect size (adjusted R2) of CRC status **(B)** and adenoma status **(C)** versus healthy controls on gut viral composition in each dataset. (**D**) The family-level composition of gut virome in each dataset. (**E** to **F)**, Comparison of relative abundance of *Siphoviridae* **(E)** and *Myoviridae* **(F)** between healthy controls and CRC patients in each dataset. Absolute z-score above 2 was considered as statistically significant. Z-score above 0 was considered as CRC-enriched, while Z-score below 0 was considered as control-enriched.

Finally, we did a compositional analysis of CRC and adenoma patients and healthy controls at the viral family level and excluded the unclassified vOTUs at the family level, which accounted for nearly 67% of total sequences. For all datasets, the gut virome was dominated by *Siphoviridae*, *Myoviridae*, *Podoviridae*, and *Quimbyviridae* (Fig. 1D). Compared with the healthy controls, *Siphoviridae* was significantly depleted in CRC patients in study Feng_2015 (*q*=0.015) and approached significant level in study Wirbel_2019 (*q*=0.087), and with meta-analysis coefficient estimate (μ)=-0.14 (*q*=0.26; Fig. 1E; Fig. S3A). *Microviridae*, a family of small ssDNA viruses, was significantly enriched in the CRC patients compared with healthy controls in 5 of 9 studies, with a meta-analysis μ=0.34 (*q*=0.02; Fig. 1F; Fig. S3A). *Autographiviridae* and *Gratiaviridae*, were also considerably enriched in the CRC patients (meta-analysis *q*<0.05; Fig. S3A). In addition, the differential family between adenoma patients and healthy controls was few, except for two clades, *Podoviridae* and *unclassified_Caudovirales*, which showed nearly significant differences between them (meta-analysis *q*<0.10; Fig. S3B).

### Identification of CRC-associated viral signatures

Given the large effect of study heterogeneity on viral shifts, we performed a more accurate meta-analysis approach to identify CRC-associated viral biomarkers across nine datasets. For each study, a Wilcoxon rank-sum test was performed on the vOTUs relative abundance profile between patients and controls. In most studies, a substantial enrichment of a set of vOTUs with very small P values was observed as compared to the expected distribution under the null hypothesis (Fig. S4A-B), indicating that some of these vOTUs are actual CRC-associated viral signatures. Based on this, we selected 516 vOTUs that had significant abundance difference (*p*<0.05 in Wilcoxon rank-sum test) and a consistent trend between patients and controls in at least 4 of 9 studies. Then, we pooled evidence of differential abundance across datasets by random effects meta-analysis and further identified 407 vOTUs at a false discovery rate (FDR) <0.01. Finally, 405 CRC-associated vOTUs were identified after adjusting for confounding variables including age, BMI, and gender (Fig. S4C; Table S3).

Over 85%, 77%, and 74.8% of the 405 CRC-associated vOTUs were independently significant in Yu_2015, Wirbel_2019, and Yachida_2019 datasets, respectively (Fig. S4D), indicating that these three studies are the primary contributors to CRC viral signatures. Inversely, only less than 20% of the CRC-associated vOTUs were independently significant within datasets Hannigan_2018 and Thomas_b_2019.

In the CRC patients, 220 of the 405 biomarkers were more abundant, while 185 of them were enriched in healthy controls. The CRC-enriched vOTUs included 76 members of *Siphoviridae*, 20 *Myoviridae*, 3 *Quimbyviridae*, and 1 *Microviridae* and 120 unclassified viruses, while the control-descending vOTUs were composed of 64 *Siphoviridae*, 25 *Myoviridae*, and 96 unclassified viruses (Fig. 2A; Table S3). We performed a host assignment of the vOTUs based on their homology or CRISPR spacers to the 4,644 prokaryotic genomes from Unified Human Gastrointestinal Genome (UHGG) collections(*27*). The analysis assigned 54.3% of the CRC-associated vOTUs into one or more prokaryotic hosts. The control-enriched vOTUs had a large proportion (23.2%) of *Ruminococcaceae* phages, whereas only 4.5% of the CRC-enriched vOTUs were those (Fisher’s exact test *q*<0.001; Fig. 2A; Table S3). Inversely, the CRC-enriched vOTUs contained significantly higher proportions of *Bacteroidaceae*, *Oscillospiraceae*, and *Peptostreptococcaceae* phages than the control-enriched viruses (Fisher’s exact test *q*<0.05). At the genus level, 37 control-enriched vOTUs were *Faecalibacterium* phages and 6 were *Roseburia* phages (Table S3). These two taxa are well-known SCFA-producers and have shown beneficial effects on multiple common disorders(*28, 29*), though they are not usually deficient in the gut microbiome of CRC patients. Several other viruses that were predicted to infect species of *Porphyromonas* (5 vOTUs), *Fusobacterium* (4 vOTUs), and *Hungatella* (3 vOTUs) were enriched in the CRC virome, in agreement with previous studies showing that these taxa are overgrown in the CRC bacteriome(*2, 3*).

**Fig. 2.**
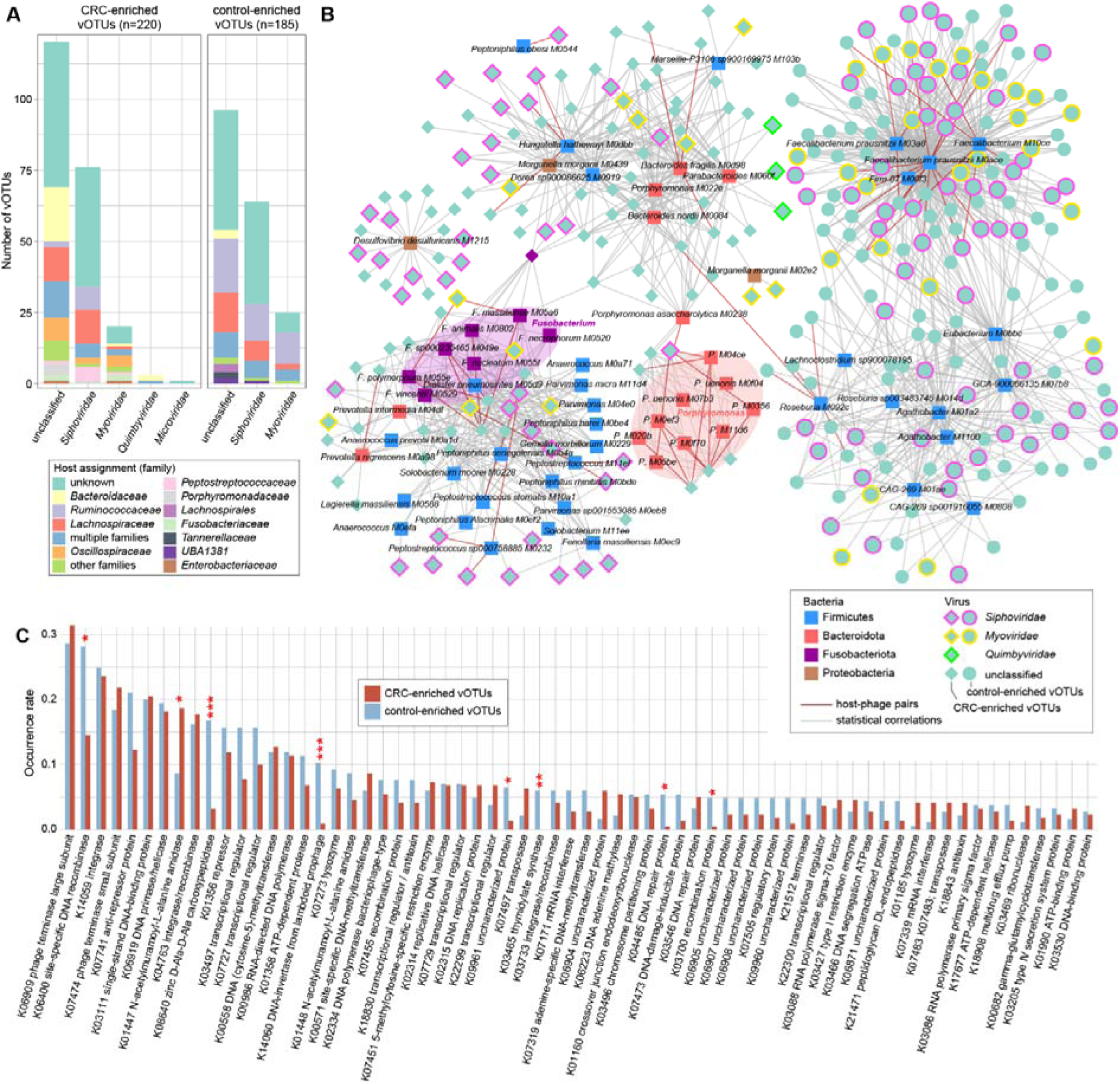
The CRC-associated viral signatures. **(A**) The family-level taxonomy and host assignment of CRC-associated vOTUs. The vOTUs are grouped at the family level, and their hosts are shown at the family level. The number of vOTUs that had more than one predicted host are colored in blue (multiple families). (**B**) The interaction network between CRC-associated viruses and bacteria. The network was constructed based on host-phage pairs and statistical co-abundance (SparCC *r*>0.35 and *q*<0.01) or co-occurrence (Fisher’s exact test, odd ratio>100 and *q*<0.01) correlations between viruses and bacteria. (**C**) The comparison of the occurrence rate of KOs detected in no fewer than 10 CRC-associated vOTUs between CRC-enriched and control-enriched vOTUs. Statistical test was performed using Fisher’s exact test, and adjusted using the FDR method. * *q* < 0.05, ** *q* < 0.01, *** *q* < 0.001.

To further investigate the interactions between CRC-associated viruses and bacteria, we identified 83 CRC-associated bacterial species from the datasets using the same approach as with virome (meta-analysis *q*<0.01; Table S4) and performed correlation analysis with 405 vOTU biomarkers. We revealed a large virus-bacterium interaction network (Fig. 2B), consisting of a total of 1,331 interactions that included 62 host-phage pairs and 1,269 statistical co-abundance or co-occurrence correlations. Diverse groups of bacteria, including the CRC-enriched *Fusobacterium* spp., *Porphyromonas* spp., *Peptoniphilus* spp., *Hungatella hathewayi*, *Bacteroides fragilis*, and *Desulfovibrio desulfuricans* and the control-enriched *Faecalibacterium prausnitzii*, *Roseburia* spp., and *Agathobacter* spp. drove the major correlations in the network (Table S4), suggesting that they may play keystone roles in the CRC gut ecosystem. The network spanned 310 vOTUs, which are potential bacterium-dependent viruses that may impact on host CRC status in cooperating with the corresponding bacteria. In addition, the remaining 85 out of 95 vOTUs might independently act on disease, as they were enriched in CRC patients (Table S3). To characterize the functional potential of the CRC-associated viruses, we annotated the proteins of 405 vOTUs using the KEGG (Kyoto Encyclopedia of Genes and Genomes) database(*30*). The KEGG orthology (KO) approach was used to analyze 19.6% (4,398/22,410) of the viral proteins that covered a total of 1,225 KOs for analysis. Statistically,14 KOs were significantly differed in frequency between the CRC-enriched and control-enriched vOTUs (Fisher’s exact test, *q*<0.05; Table S5). Several enzymes, including K08640 (zinc D-Ala-D-Ala carboxypeptidase), K14060 (DNA-invertase from lambdoid prophage), K03465 (thymidylate synthase), K06400 (site-specific DNA recombinase), K04485 (DNA repair protein), and K03700 (recombination protein) were more frequently encoded in the control group-enriched viruses, whereas K01447 (N-acetylmuramoyl-L-alanine amidase) was more abundant in the CRC-enriched viruses (Fig. 2C).

### Constructing the CRC predictive model using viral signatures

To detect the diagnostic ability of gut viral signatures in CRC, we performed intra-dataset and cross-dataset prediction and validation on the overall set of 554 CRC and 546 control samples based on their relative abundances of 405 vOTUs. Two machine learning algorithms, least absolute shrinkage and selection operator (LASSO) and random forest (RF), were used for modeling and testing (see Methods). In intra-dataset, we observed performances ranging in area under the receiver operating characteristic curve (AUC) score from 0.582 to 0.900 (average 0.779) for the LASSO algorithm and from 0.631 to 0.870 (average 0.752) for the RF algorithm (Fig. 3A-B). The Hannigan_2018 study obtained the lowest AUCs in both algorithms, which could potentially be explained by its small sample size. In cross-dataset prediction and validation, we obtained pairwise AUCs ranging from 0.539 to 0.926 (average 0.753) for LASSO algorithm and from 0.539 to 0.890 (average 0.737) for the RF algorithm.

**Fig. 3.**
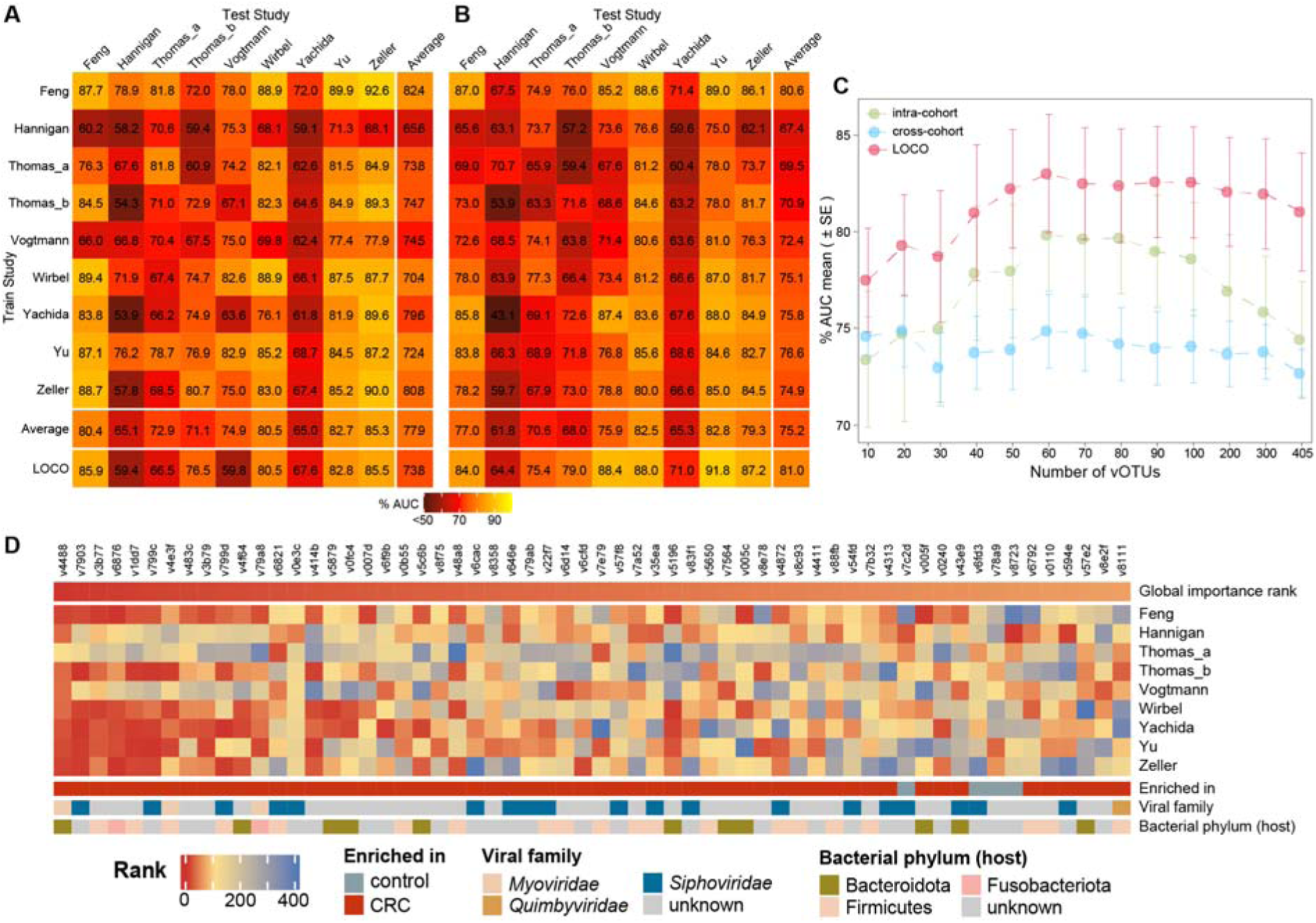
The prediction model of CRC status based on relative abundances of 405 CRC-associated vOTUs. **(A** to **B**) Performance assessment as AUC scores of intra-dataset and cross-dataset predictions using least absolute shrinkage and selection operator **(A)** and random forest models **(B)** in predicting CRC status. The model of intra-dataset prediction (diagonal) was validated using five repeats of five-fold cross-validations. The model of cross-dataset prediction (off-diagonal) was built on the dataset corresponding to each row and validated on the dataset corresponding to each column. The LOCO row refers to leave-one-cohort-out (LOCO) analysis in which models were built on eight datasets combined and validated on the remaining one corresponding to each column. (**C**) Average AUC values for different numbers of CRC-associated vOTUs using random forest models. (**D**) the heat map showing the 60 most important ranking vOTUs in the random forest model. The global importance rank refers to the mean rank of each vOTU for all studies. The feature importance was calculated by the “mean decrease accuracy” method.

To overcome the sample size limitations on single studies, we performed a leave-one-cohort-out (LOCO) analysis, in which models were built on eight datasets combined and validated on the left-out dataset, for each dataset in turn. The LOCO AUCs ranged from 0.594 to 0.859 (average 0.779) and from 0.644 to 0.918 (average 0.810) for nine studies using the LASSO and RF models, respectively (Fig. 3A-B). In addition, adding the host properties (i.e., age, BMI, and gender) didn’t improve the predicting performance for both algorithms (Fig. S5). Taken together, our intra-dataset, cross-dataset, and combined-dataset analyses suggest that the gut viral signatures are efficient for distinguishing CRC patients from healthy controls.

Finally, to generate a minimal set of vOTU signatures, we calculated the global importance ranks of 405 vOTUs from the RF models of all studies. Using the RF algorithm, CRC was accurately identified with an average cross-dataset AUC 0.798 and LOCO AUC 0.830 when using a subset of 60 top importance vOTUs (Fig. 3C; Fig. S6). Notably, 56 of the top importance vOTUs were CRC-enriched biomarkers, which included 4 *Peptostreptococcus*, 3 *Fusobacterium*, and 3 *Porphyromonas* phages (Fig. 3D; Table S3).

### Validation of CRC viral markers in independent cohorts

To validate the efficiency of CRC viral signatures in the independent cohort, we recruited 27 CRC patients having matched age, BMI and gender and healthy controls and performed whole-metagenome shotgun sequencing of their fecal samples (Table S6). In this new cohort, 207 out of the 405 CRC-associated vOTUs, including 46 CRC-enriched and 161 control-enriched vOTUs, were significantly differed in relative abundance between patients and controls with a consistent trend with the meta-analysis of nine studies (*p*<0.05 in Wilcoxon rank-sum test; Fig. 4A; Table S3). Compared with the control-enriched vOTUs, the lower rate of CRC-enriched vOTUs might be contributed by their lower occurrence rate in fecal samples of the new cohort, as well as in samples of the above nine studies (Fig. S7). Using the RF algorithm, we trained two models based on the abundances of 405 vOTUs and 60 top importance vOTUs in the original nine cohorts. These models obtained the AUC of 0.906 and 0.877, respectively, in distinguishing between CRC and controls in the independent new cohort (Fig. 4B). We also validated the efficiency of CRC viral signatures in the recently published cohort of 100 onset CRC patients and 100 healthy controls (Yang_2021)(*7*). The models obtained an acceptable diagnostic efficacy in distinguishing onset CRC patients and controls (Fig. 4D). These findings suggest high repeatability and predictive power of CRC viral markers in independent cohorts.

**Fig. 4.**
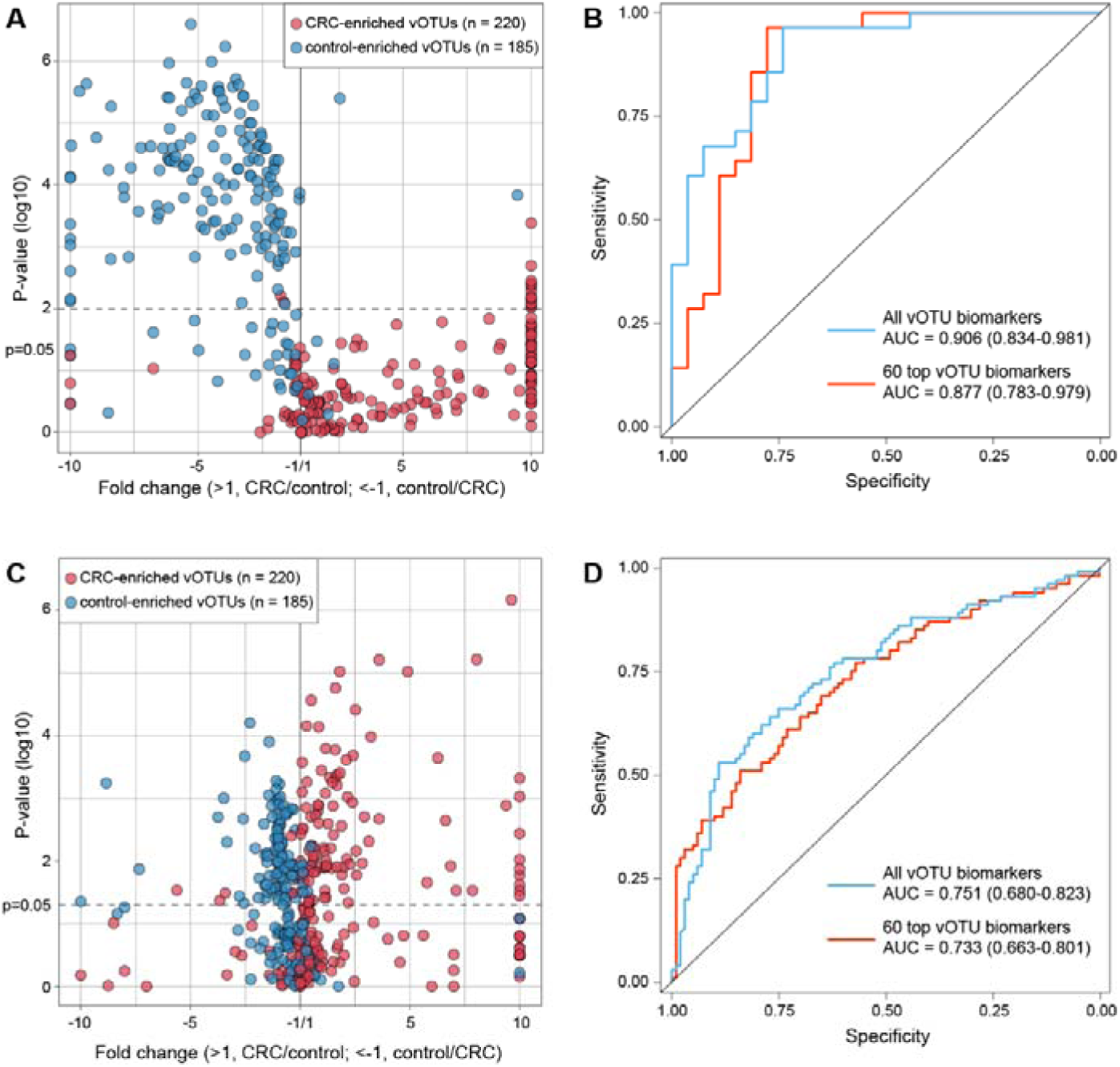
Validation of 405 CRC-associated vOTUs in independent cohorts. **(A** and **C)** Volcano plots showing fold change and statistical significance in CRC-associated vOTUs abundance between CRC patients and healthy controls recruited in this study **(A)** and Yang’s study **(C)**. (**B** and **D)** Performance assessment as AUC scores for CRC-associated vOTUs using random forest models in the cohort from this study **(B)** and Yang’s study **(D)**, respectively. The model was built by combining nine published datasets and was validated by independent cohorts. Blue curve, the model based on all CRC-associated vOTUs abundances. Red curve, the model based on 60 most important ranking vOTUs abundances.

### Gut viral signatures in adenoma

Lastly, we profiled the gut viral profiles of adenoma patients (n = 182) from five studies and compared them with those of corresponding healthy controls (n = 388; Table 1) to explore adenoma viral signatures. A meta-analysis based on the aforementioned approach identified 88 significant adenoma-associated vOTUs (meta-analysis *q*<0.05; Table S7). In adenoma patients, 47 of these vOTUs were more abundant, including 20 Siphoviridae, 3 Myoviridae, a Quimbyviridae, and a crAss-like viruses; while 41 vOTUs were enriched in controls, including 20 *Siphoviridae*, 3 *Myoviridae*, and 1 *Flandersviridae* viruses. Host assignment showed that the adenoma-enriched vOTUs included 4 *Enterobacteriaceae*, 3 *Bacteroidaceae*, and 3 *Oscillospiraceae* phages, whereas 3 of control-enriched vOTUs were *Bifidobacteriaceae* phages (Table S7). Additionally, adenoma and CRC shared 4 differential viral markers, including 2 unclassified adenoma/CRC-enriched and 2 control-enriched vOTUs that belonged to *Siphoviridae*.

We performed machine learning for adenoma prediction and observed the intra-dataset, cross-dataset, and LOCO average AUCs of 0.700, 0.686, and 0.772, respectively. In five adenoma studies using the LASSO algorithm (Fig. 5a), average AUCs of 0.690, 0.698, and 0.760 using the RF algorithm (Fig. 5b), suggesting the validity of adenoma-specific viral markers.

**Fig. 5.**
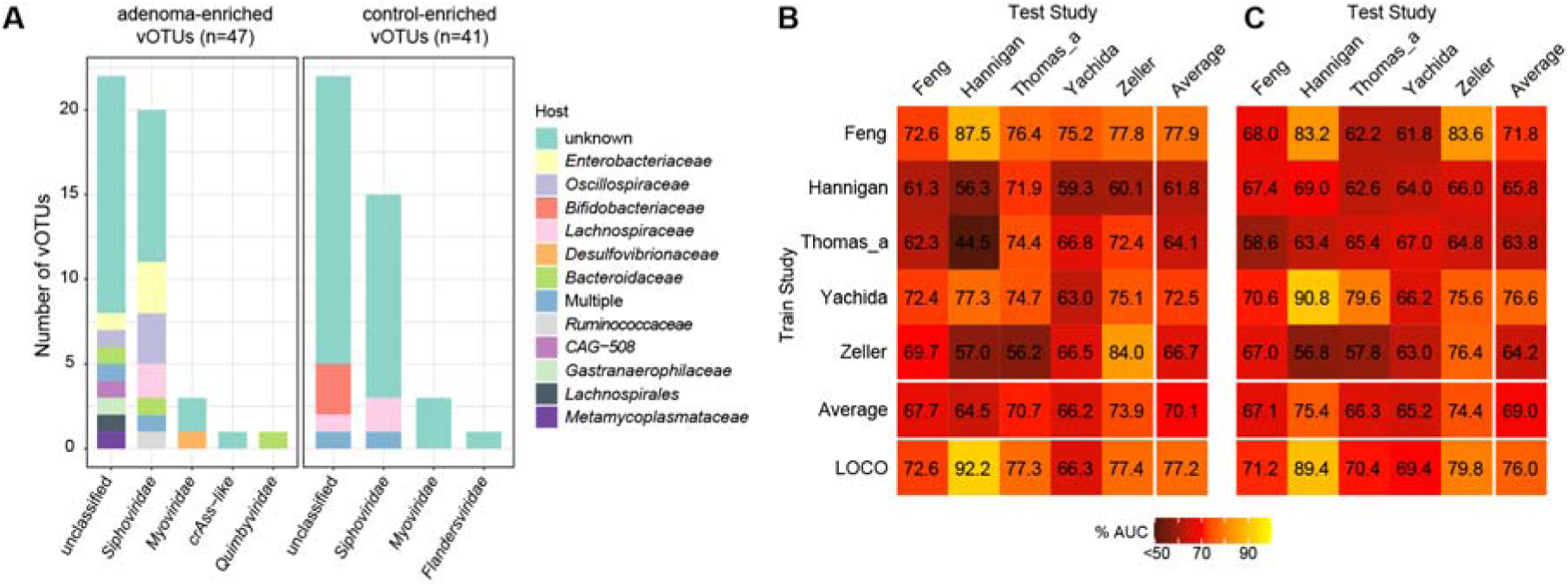
The adenoma-associated viral signatures. (**A)** The family-level taxonomy and host assignment of adenoma-associated vOTUs. The vOTUs were grouped at the family level, and their hosts are shown at the family level. The number of vOTUs that has more than one predicted hosts are colored in blue (multiple families). (**B** to **C**) Performance assessment as AUC scores of intra-dataset and cross-dataset predictions using LASSO **(B)** and random forest **(C)** model in predicting adenoma status. The model of intra-cohort prediction (diagonal) was validated using five repeats of 5-fold cross-validations. The model of cross-cohort prediction (off-diagonal) was built on the dataset corresponding to each row and validated on the dataset corresponding to each column. The LOCO row refers to leave-one-cohort-out (LOCO) analysis in which models were built on eight datasets combined and validated on the remaining one corresponding to each column.

## Discussion

Although a series of studies indicated intestinal dysbacteriosis in CRC patients, most of them focused on the characterization of the gut bacteriome(*2-4, 7, 31*), and only a few studies explored a limited number of known viruses(*16, 17*). In this study, we introduced a more comprehensive database of viral genomes that contained 37,030 non-redundant viral genomes derived from the *de novo* assembly of 1,282 fecal metagenomes from 9 published cohorts. Over half of the genomes in this database were undetected by the current available viral databases, which supported that our database was necessary to gain insight into the alteration of gut viral communities in CRC patients. On the basis of this database, we found that the gut virome diversities of CRC patients were comparable to those of healthy controls in most of the cohorts collected in this study, consistent with the finding of previous studies(*16, 18*). The viral diversity parameters of samples were highly consistent with their bacterial diversities, which might be explained by the fact that most of the viruses in this study resided within bacterial cells.

We identified 405 reproducible vOTUs associated with CRC using meta-analysis based on 9 independent cohorts. The majority of these vOTUs are classified as viral dark matter because of the absence of referral genomes. We found that most of these vOTUs (310/405) were closely connected with bacterial biomarkers for CRC, and they aggregated into a multi-centric virus-bacteria interaction network (Fig. 2B). The bacterial biomarkers were central members within the network, which suggested that these viruses might indirectly influence CRC development by altering bacterial biomarkers. Among this network, the central members of two hubs were mainly the butyrate-producing bacteria from the genera *Faecalibacterium*, *Roseburia*, and *Agathobacter*. Some studies implied that butyrate producers, *Faecalibacterium prausnitzii* in particular, were potential probiotics with anti-tumorigenic properties and might contribute to preventing CRC development(*7, 32*). Our results showed that a group of viruses, reduced in CRC patients, were closely connected with butyrate-producing bacteria, which suggested these viruses may contribute to tumorigenesis by modulating the butyrate-producing bacteria in human gut. Another hub in this network involved a variety of common oral bacteria from the genera *Anaerococcus*, *Fusobacterium*, *Peptostreptococcus*, *Parvimonas*, and *Solobacterium*. Many studies have reported that these bacteria are significantly enriched in fecal samples from CRC patients(*2, 4*). Translocation of oral microbes to the gut caused an increased abundance of them in cancer patients. Particularly, we observed that one N-acetylmuramoyl-L-alanine amidase (K01447) was more abundant in the CRC-enriched viruses than in CON-enriched viruses, indicating that it can digest the peptidoglycan of the bacterial cell wall and disrupt the biofilms(*33, 34*). CRC-enriched viruses, coexisting with bacterial biomarkers commonly present in the oral cavity, might affect CRC development by disrupting bacterial biofilms to promote dispersal of these bacteria. Overall, our results partly explained and put a spotlight on how CRC-associated viruses can influence cancer progression. However, our findings are data-driven, and further studies will be necessary to uncover the linkage between viruses, bacteria, and carcinogenesis by in vitro and in vivo experimental validation.

Furthermore, gut bacteria-based CRC prediction models have demonstrated high diagnostic potential; however, the models are not available in all cohorts(*2, 3*), implying that the identification of new potential predictors was required. Although gut viruses have not been regarded as the effective CRC predictors for a long time(*16, 17*), a recent study reported the potential of gut viruses for the classifiability of controls versus patients with CRC(*2*). In this study, we performed viral biomarker-based random forest models with LOCO validation and were able to distinguish CRC patients from healthy controls with an average performance of 0.81 AUC which was comparable to the performance of models based on bacterial biomarkers (Fig. 3B). Importantly, we also validated the efficiency of CRC viral signatures in two independent cohorts. For the newly-recruited cohort in this study, CRC viral signatures displayed excellent potential for classifiability of controls versus patients (Fig. 4B). For other independent cohorts, the models only obtained acceptable diagnostic efficacy in distinguishing onset CRC patients and controls (Fig. 4D). Alteration of certain CRC viral signatures is induced by therapeutic approaches. Among CRC viral signatures, a phage (v4488) infecting *Bacteroides xylanisolvens* was ranked as the top contributor to CRC prediction models (Fig. 3D). But *Bacteroides xylanisolvens* is not reported as the CRC biomarker so far, which suggests that certain gut phages play an irreplaceable role in CRC prediction. In addition, several top-ranking important biomarkers infected the CRC-associated species of the genera including *Peptostreptococcus*, *Fusobacterium*, and *Porphyromonas* (Fig. 3D), suggesting that these viruses might be the CRC predictors with similar performances as their hosts. Taken together, our findings showed that viral biomarkers would be the new efficient predictors for CRC diagnosis.

Moreover, colorectal adenomas are regarded as precursor lesions of CRC. The detection of colorectal adenomas could reduce the risk of CRC and improve survival rates. Therefore, we also identified a series of adenomas-associated viruses and assessed the performance of prediction models for adenoma status. Both LASSO and RF algorithms with LOCO approaches demonstrated that adenomas patients and healthy controls could be distinguished with acceptable accuracy (average AUC > 0.75). Similar results were observed in the bacteria-based prediction models for adenoma status(*31*). These findings indicate the dysbiosis of intestinal viral communities in colorectal adenomas patients, highlighting the potential role of viral biomarkers in the diagnosis of colorectal adenomas.

In conclusion, we comprehensively observed shifts in intestinal viral composition in CRC patients and also described interactions between viral biomarkers and bacteria. Although the precise mechanisms of how these viruses cause tumorigenesis are still unclear, our works provide several potential explanations. Furthermore, the present findings indicate that gut viromes could be used to improve microbiome-based CRC diagnostics. Our analysis strongly suggested that more research is needed to determine the precise role of the omitted gut virome in CRC development.

## Conclusion

This work demonstrates the differences of gut virome between the healthy, adenoma patients and CRC patients and obtains clinically meaningful viral markers based on statistical analysis. It provides a new direction for the early diagnosis and pathological research of adenoma and CRC in the future.

## Acknowledgments

This work was financially supported by National Natural Science Foundation of China (No. 81930112 and No. 81902037), Distinguished professor of Liaoning Province (XLYC2002008), and Dalian Science and Technology Leading Talents Project (2019RD15).

## Data and materials availability

All data needed to evaluate the conclusions in the paper are present in the paper and/or the Supplementary Materials.

## Reference

1. M. Arnold, M. S. Sierra, M. Laversanne, I. Soerjomataram, A. Jemal, F. Bray, Global patterns and trends in colorectal cancer incidence and mortality. Gut 66, 683–691 (2017).

2. A. M. Thomas, P. Manghi, F. Asnicar, E. Pasolli, F. Armanini, M. Zolfo, F. Beghini, S. Manara, N. Karcher, C. Pozzi, S. Gandini, D. Serrano, S. Tarallo, A. Francavilla, G. Gallo, M. Trompetto, G. Ferrero, S. Mizutani, H. Shiroma, S. Shiba, T. Shibata, S. Yachida, T. Yamada, J. Wirbel, P. Schrotz-King, C. M. Ulrich, H. Brenner, M. Arumugam, P. Bork, G. Zeller, F. Cordero, E. Dias-Neto, J. C. Setubal, A. Tett, B. Pardini, M. Rescigno, L. Waldron, A. Naccarati, N. Segata, Metagenomic analysis of colorectal cancer datasets identifies cross-cohort microbial diagnostic signatures and a link with choline degradation. Nat Med 25, 667–678 (2019).

3. J. Wirbel, P. T. Pyl, E. Kartal, K. Zych, A. Kashani, A. Milanese, J. S. Fleck, A. Y. Voigt, A. Palleja, R. Ponnudurai, S. Sunagawa, L. P. Coelho, P. Schrotz-King, E. Vogtmann, N. Habermann, E. Nimeus, A. M. Thomas, P. Manghi, S. Gandini, D. Serrano, S. Mizutani, H. Shiroma, S. Shiba, T. Shibata, S. Yachida, T. Yamada, L. Waldron, A. Naccarati, N. Segata, R. Sinha, C. M. Ulrich, H. Brenner, M. Arumugam, P. Bork, G. Zeller, Meta-analysis of fecal metagenomes reveals global microbial signatures that are specific for colorectal cancer. Nat Med 25, 679–689 (2019).

4. S. Yachida, S. Mizutani, H. Shiroma, S. Shiba, T. Nakajima, T. Sakamoto, H. Watanabe, K. Masuda, Y. Nishimoto, M. Kubo, F. Hosoda, H. Rokutan, M. Matsumoto, H. Takamaru, M. Yamada, T. Matsuda, M. Iwasaki, T. Yamaji, T. Yachida, T. Soga, K. Kurokawa, A. Toyoda, Y. Ogura, T. Hayashi, M. Hatakeyama, H. Nakagama, Y. Saito, S. Fukuda, T. Shibata, T. Yamada, Metagenomic and metabolomic analyses reveal distinct stage-specific phenotypes of the gut microbiota in colorectal cancer. Nat Med 25, 968–976 (2019).

5. A. D. Kostic, E. Chun, L. Robertson, J. N. Glickman, C. A. Gallini, M. Michaud, T. E. Clancy, D. C. Chung, P. Lochhead, G. L. Hold, E. M. El-Omar, D. Brenner, C. S. Fuchs, M. Meyerson, W. S. Garrett, Fusobacterium nucleatum potentiates intestinal tumorigenesis and modulates the tumor-immune microenvironment. Cell host & microbe 14, 207–215 (2013).

6. J. Hong, F. Guo, S.-Y. Lu, C. Shen, D. Ma, X. Zhang, Y. Xie, T. Yan, T. Yu, T. Sun, Y. Qian, M. Zhong, J. Chen, Y. Peng, C. Wang, X. Zhou, J. Liu, Q. Liu, X. Ma, Y.-X. Chen, H. Chen, J.-Y. Fang, F. nucleatum targets lncRNA ENO1-IT1 to promote glycolysis and oncogenesis in colorectal cancer. Gut, gutjnl-2020-322780 (2020).

7. Y. Yang, L. Du, D. Shi, C. Kong, J. Liu, G. Liu, X. Li, Y. Ma, Dysbiosis of human gut microbiome in young-onset colorectal cancer. Nature communications 12, 6757 (2021).

8. R. Guo, S. Li, Y. Zhang, Y. Zhang, G. Wang, Y. Ma, Q. Yan, Dysbiotic oral and gut viromes in untreated and treated rheumatoid arthritis patients. bioRxiv, (2021).

9. A. G. Clooney, T. D. S. Sutton, A. N. Shkoporov, R. K. Holohan, K. M. Daly, O. O’Regan, F. J. Ryan, L. A. Draper, S. E. Plevy, R. P. Ross, C. Hill, Whole-Virome Analysis Sheds Light on Viral Dark Matter in Inflammatory Bowel Disease. Cell Host Microbe 26, 764–778 e765 (2019).

10. X. Zhang, D. Zhang, H. Jia, Q. Feng, D. Wang, D. Liang, X. Wu, J. Li, L. Tang, Y. Li, Z. Lan, B. Chen, Y. Li, H. Zhong, H. Xie, Z. Jie, W. Chen, S. Tang, X. Xu, X. Wang, X. Cai, S. Liu, Y. Xia, J. Li, X. Qiao, J. Y. Al-Aama, H. Chen, L. Wang, Q. J. Wu, F. Zhang, W. Zheng, Y. Li, M. Zhang, G. Luo, W. Xue, L. Xiao, J. Li, W. Chen, X. Xu, Y. Yin, H. Yang, J. Wang, K. Kristiansen, L. Liu, T. Li, Q. Huang, Y. Li, J. Wang, The oral and gut microbiomes are perturbed in rheumatoid arthritis and partly normalized after treatment. Nat Med 21, 895–905 (2015).

11. E. A. Franzosa, A. Sirota-Madi, J. Avila-Pacheco, N. Fornelos, H. J. Haiser, S. Reinker, T. Vatanen, A. B. Hall, H. Mallick, L. J. McIver, J. S. Sauk, R. G. Wilson, B. W. Stevens, J. M. Scott, K. Pierce, A. A. Deik, K. Bullock, F. Imhann, J. A. Porter, A. Zhernakova, J. Fu, R. K. Weersma, C. Wijmenga, C. B. Clish, H. Vlamakis, C. Huttenhower, R. J. Xavier, Gut microbiome structure and metabolic activity in inflammatory bowel disease. Nature microbiology 4, 293–305 (2019).

12. D. C. Damin, P. K. Ziegelmann, A. P. Damin, Human papillomavirus infection and colorectal cancer risk: a meta-analysis. Colorectal Dis 15, e420–428 (2013).

13. S. Bedri, A. A. Sultan, M. Alkhalaf, A. E. Al Moustafa, S. Vranic, Epstein-Barr virus (EBV) status in colorectal cancer: a mini review. Hum Vaccin Immunother 15, 603–610 (2019).

14. F. H. Su, T. N. Le, C. H. Muo, S. A. Te, F. C. Sung, C. C. Yeh, Chronic Hepatitis B Virus Infection Associated with Increased Colorectal Cancer Risk in Taiwanese Population. Viruses 12, (2020).

15. H. P. Chen, J. K. Jiang, P. Y. Lai, C. Y. Chen, T. Y. Chou, Y. C. Chen, C. H. Chan, S. F. Lin, C. Y. Yang, C. Y. Chen, C. H. Lin, J. K. Lin, D. M. Ho, W. L. Cho, Y. J. Chan, Tumoral presence of human cytomegalovirus is associated with shorter disease-free survival in elderly patients with colorectal cancer and higher levels of intratumoral interleukin-17. Clin Microbiol Infect 20, 664–671 (2014).

16. G. D. Hannigan, M. B. Duhaime, M. T. t. Ruffin, C. C. Koumpouras, P. D. Schloss, Diagnostic Potential and Interactive Dynamics of the Colorectal Cancer Virome. mBio 9, (2018).

17. G. Nakatsu, H. Zhou, W. K. K. Wu, S. H. Wong, O. O. Coker, Z. Dai, X. Li, C. H. Szeto, N. Sugimura, T. Y. Lam, A. C. Yu, X. Wang, Z. Chen, M. C. Wong, S. C. Ng, M. T. V. Chan, P. K. S. Chan, F. K. L. Chan, J. J. Sung, J. Yu, Alterations in Enteric Virome Are Associated With Colorectal Cancer and Survival Outcomes. Gastroenterology 155, 529–541 e525 (2018).

18. S. Shen, D. Huo, C. Ma, S. Jiang, J. Zhang, Expanding the Colorectal Cancer Biomarkers Based on the Human Gut Phageome. Microbiol Spectr, e0009021 (2021).

19. G. Zeller, J. Tap, A. Y. Voigt, S. Sunagawa, J. R. Kultima, P. I. Costea, A. Amiot, J. Bohm, F. Brunetti, N. Habermann, R. Hercog, M. Koch, A. Luciani, D. R. Mende, M. A. Schneider, P. Schrotz-King, C. Tournigand, J. Tran Van Nhieu, T. Yamada, J. Zimmermann, V. Benes, M. Kloor, C. M. Ulrich, M. von Knebel Doeberitz, I. Sobhani, P. Bork, Potential of fecal microbiota for early-stage detection of colorectal cancer. Mol Syst Biol 10, 766 (2014).

20. J. Yu, Q. Feng, S. H. Wong, D. Zhang, Q. Y. Liang, Y. Qin, L. Tang, H. Zhao, J. Stenvang, Y. Li, X. Wang, X. Xu, N. Chen, W. K. Wu, J. Al-Aama, H. J. Nielsen, P. Kiilerich, B. A. Jensen, T. O. Yau, Z. Lan, H. Jia, J. Li, L. Xiao, T. Y. Lam, S. C. Ng, A. S. Cheng, V. W. Wong, F. K. Chan, X. Xu, H. Yang, L. Madsen, C. Datz, H. Tilg, J. Wang, N. Brunner, K. Kristiansen, M. Arumugam, J. J. Sung, J. Wang, Metagenomic analysis of faecal microbiome as a tool towards targeted non-invasive biomarkers for colorectal cancer. Gut 66, 70–78 (2017).

21. Q. Feng, S. Liang, H. Jia, A. Stadlmayr, L. Tang, Z. Lan, D. Zhang, H. Xia, X. Xu, Z. Jie, L. Su, X. Li, X. Li, J. Li, L. Xiao, U. Huber-Schonauer, D. Niederseer, X. Xu, J. Y. Al-Aama, H. Yang, J. Wang, K. Kristiansen, M. Arumugam, H. Tilg, C. Datz, J. Wang, Gut microbiome development along the colorectal adenoma-carcinoma sequence. Nat Commun 6, 6528 (2015).

22. E. Vogtmann, X. Hua, G. Zeller, S. Sunagawa, A. Y. Voigt, R. Hercog, J. J. Goedert, J. Shi, P. Bork, R. Sinha, Colorectal Cancer and the Human Gut Microbiome: Reproducibility with Whole-Genome Shotgun Sequencing. PLoS One 11, e0155362 (2016).

23. A. C. Gregory, O. Zablocki, A. A. Zayed, A. Howell, B. Bolduc, M. B. Sullivan, The Gut Virome Database Reveals Age-Dependent Patterns of Virome Diversity in the Human Gut. Cell Host Microbe 28, 724–740 e728 (2020).

24. L. F. Camarillo-Guerrero, A. Almeida, G. Rangel-Pineros, R. D. Finn, T. D. Lawley, Massive expansion of human gut bacteriophage diversity. Cell 184, 1098–1109 e1099 (2021).

25. S. Nayfach, A. P. Camargo, F. Schulz, E. Eloe-Fadrosh, S. Roux, N. C. Kyrpides, CheckV assesses the quality and completeness of metagenome-assembled viral genomes. Nat Biotechnol, (2020).

26. S. Nayfach, D. Paez-Espino, L. Call, S. J. Low, H. Sberro, N. N. Ivanova, A. D. Proal, M. A. Fischbach, A. S. Bhatt, P. Hugenholtz, N. C. Kyrpides, Metagenomic compendium of 189,680 DNA viruses from the human gut microbiome. Nat Microbiol 6, 960–970 (2021).

27. A. Almeida, S. Nayfach, M. Boland, F. Strozzi, M. Beracochea, Z. J. Shi, K. S. Pollard, E. Sakharova, D. H. Parks, P. Hugenholtz, N. Segata, N. C. Kyrpides, R. D. Finn, A unified catalog of 204,938 reference genomes from the human gut microbiome. Nature biotechnology 39, 105–114 (2021).

28. K. Machiels, M. Joossens, J. Sabino, V. De Preter, I. Arijs, V. Eeckhaut, V. Ballet, K. Claes, F. Van Immerseel, K. Verbeke, M. Ferrante, J. Verhaegen, P. Rutgeerts, S. Vermeire, A decrease of the butyrate-producing species Roseburia hominis and Faecalibacterium prausnitzii defines dysbiosis in patients with ulcerative colitis. Gut 63, 1275–1283 (2014).

29. L. Zhou, M. Zhang, Y. Wang, R. G. Dorfman, H. Liu, T. Yu, X. Chen, D. Tang, L. Xu, Y. Yin, Y. Pan, Q. Zhou, Y. Zhou, C. Yu, Faecalibacterium prausnitzii Produces Butyrate to Maintain Th17/Treg Balance and to Ameliorate Colorectal Colitis by Inhibiting Histone Deacetylase 1. Inflamm Bowel Dis 24, 1926–1940 (2018).

30. M. Kanehisa, M. Furumichi, M. Tanabe, Y. Sato, K. Morishima, KEGG: new perspectives on genomes, pathways, diseases and drugs. Nucleic Acids Res 45, D353–D361 (2017).

31. Y. Wu, N. Jiao, R. Zhu, Y. Zhang, D. Wu, A. J. Wang, S. Fang, L. Tao, Y. Li, S. Cheng, X. He, P. Lan, C. Tian, N. N. Liu, L. Zhu, Identification of microbial markers across populations in early detection of colorectal cancer. Nature communications 12, 3063 (2021).

32. J. Chen, L. Vitetta, Inflammation-modulating effect of butyrate in the prevention of colon cancer by dietary fiber. Clinical colorectal cancer 17, e541–e544 (2018).

33. M. Domenech, E. García, M. Moscoso, In vitro destruction of Streptococcus pneumoniae biofilms with bacterial and phage peptidoglycan hydrolases. Antimicrobial agents and chemotherapy 55, 4144–4148 (2011).

34. A. Vermassen, S. Leroy, R. Talon, C. Provot, M. Popowska, M. Desvaux, Cell wall hydrolases in bacteria: insight on the diversity of cell wall amidases, glycosidases and peptidases toward peptidoglycan. Frontiers in microbiology 10, 331 (2019).

35. S. Chen, Y. Zhou, Y. Chen, J. Gu, fastp: an ultra-fast all-in-one FASTQ preprocessor. Bioinformatics 34, i884–i890 (2018).

36. B. Langmead, S. L. Salzberg, Fast gapped-read alignment with Bowtie 2. Nature methods 9, 357–359 (2012).

37. D. Li, C. M. Liu, R. Luo, K. Sadakane, T. W. Lam, MEGAHIT: an ultra-fast single-node solution for large and complex metagenomics assembly via succinct de Bruijn graph. Bioinformatics 31, 1674–1676 (2015).

38. S. Li, R. Guo, Y. Zhang, P. Li, F. Chen, X. Wang, J. Li, Z. Jie, Q. Lv, H. Jin, A Catalogue of 48,425 Nonredundant Viruses From Oral Metagenomes Expands the Horizon of the Human Oral Virome. under review 48.

39. J. Friedman, E. J. Alm, Inferring correlation networks from genomic survey data. PLoS Comput Biol 8, e1002687 (2012).

40. S. C. Watts, S. C. Ritchie, M. Inouye, K. E. Holt, FastSpar: rapid and scalable correlation estimation for compositional data. Bioinformatics 35, 1064–1066 (2019).

41. P. Shannon, A. Markiel, O. Ozier, N. S. Baliga, J. T. Wang, D. Ramage, N. Amin, B. Schwikowski, T. Ideker, Cytoscape: a software environment for integrated models of biomolecular interaction networks. Genome research 13, 2498–2504 (2003).

42. J. Wirbel, K. Zych, M. Essex, N. Karcher, E. Kartal, G. Salazar, P. Bork, S. Sunagawa, G. Zeller, Microbiome meta-analysis and cross-disease comparison enabled by the SIAMCAT machine learning toolbox. Genome biology 22, 1–27 (2021).

