## Supplemental Figure for "Meta-analysis of fecal metagenomes reveals global viral signatures and its diagnostic potential for colorectal cancer and adenoma"

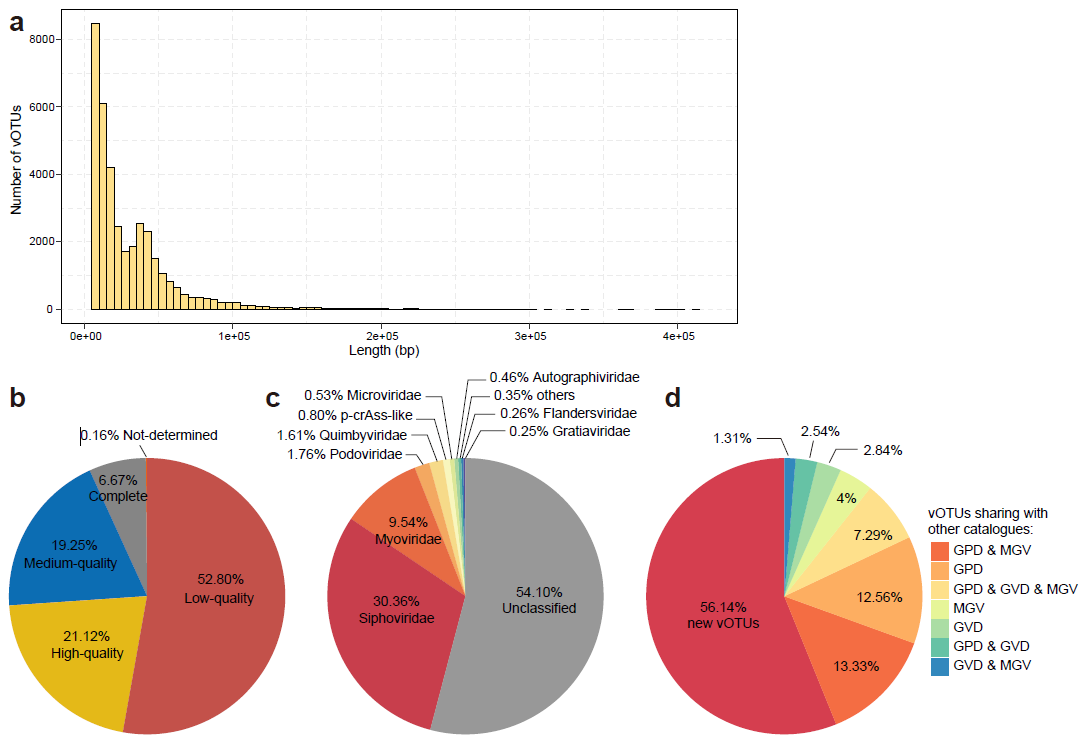

**Supplementary Figure 1. Characteristics of the nonredundant viral catalog. a**, Distribution of sequence length of 37,030 nonredundant vOTUs. **b-c**, The estimated completeness **(b)** and family-level taxonomic assignment **(c)** of vOTUs. **d**, Overlap of viruses between the nonredundant viral catalog of this study and other published viral catalogs.

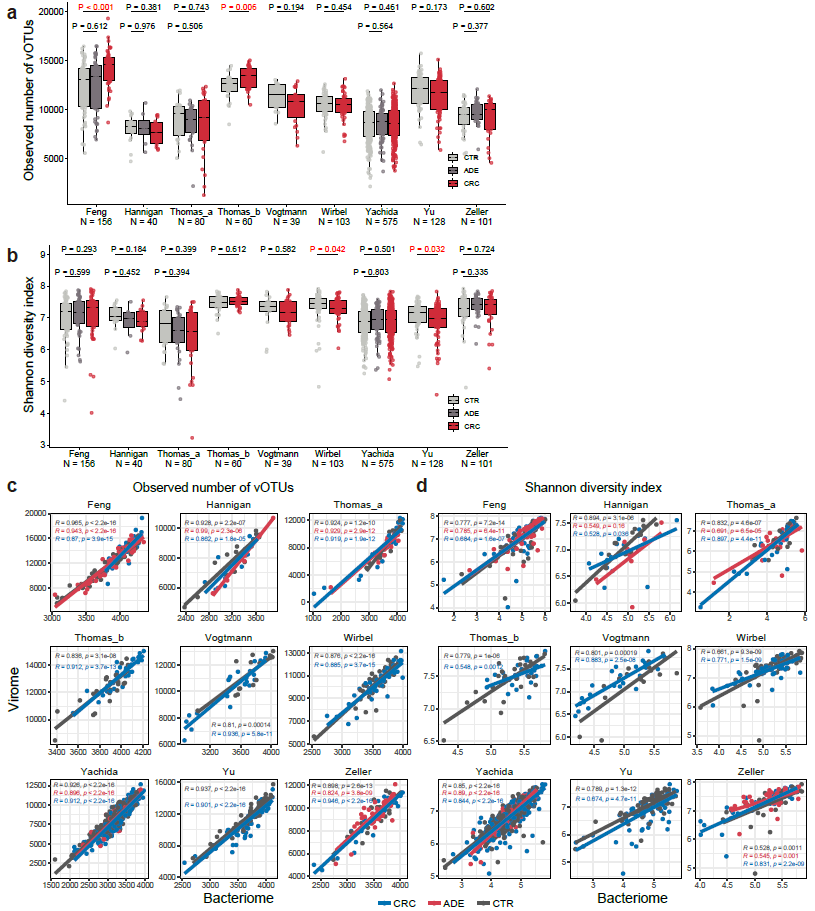

**Figure S2. Species richness and evenness of gut microbiome across CRC datasets. a**, Comparison of the observed number of vOTUs among three groups in each dataset. The significance test was performed by the two-tailed Wilcoxon rank-sum test. CTR, healthy controls; ADE, adenoma patients; CRC, colorectal carcinoma patients. **b**, Comparison of Shannon diversity index among three groups in each dataset. **c**, Scatter plot of the observed number of vOTUs of virome and bacteriome in each dataset. The correlation of alpha diversity between virome and bacteriome was visualized by a linear regression line. Correlation test was performed by the Pearson rank correlation method. **d**, Scatter plot of Shannon diversity index of virome and bacteriome in each dataset.

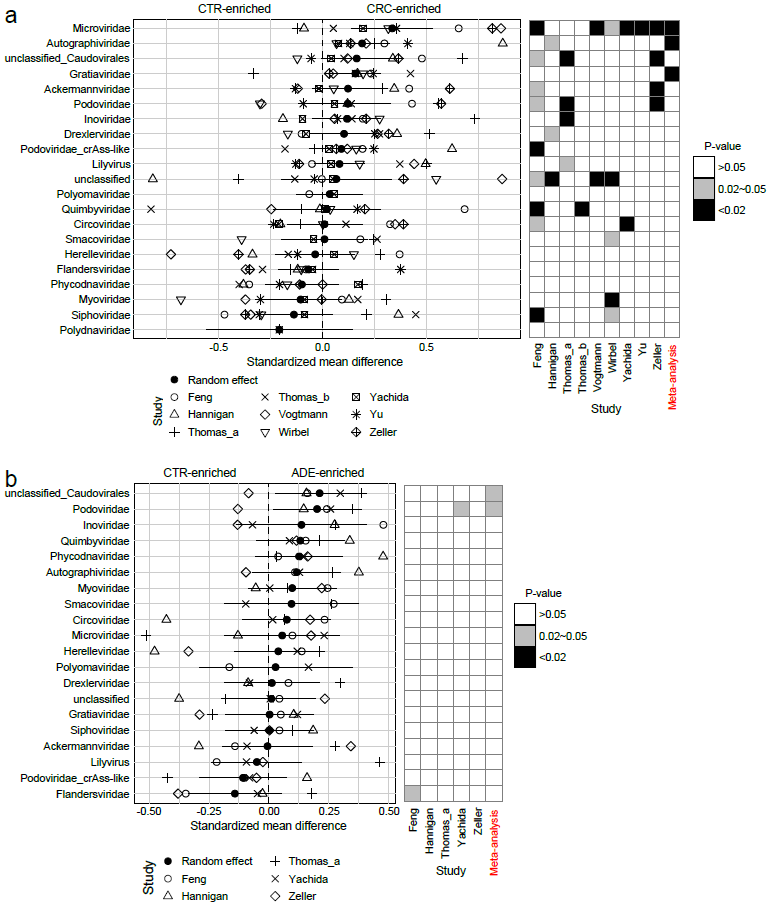

**Figure S3. Meta-analysis of the relative abundances** **of viral families in healthy controls and patients with colorectal carcinoma or adenoma.** Left panel; forest plot created using Hedges' g standardized mean differences and the random effects model based on the arcsine-square root-transformed abundances of viral families. Solid black lines indicate the 95% confidence intervals. Right panel; heat map showing abundance change significance (p-value) of each vial family between healthy controls and patients with colorectal carcinoma **(a)** or adenoma **(b)** that was estimated by meta-analysis or Wilcoxon signed-rank test in each data set. CTR, healthy controls; ADE, adenoma patients; CRC, colorectal carcinoma patients.

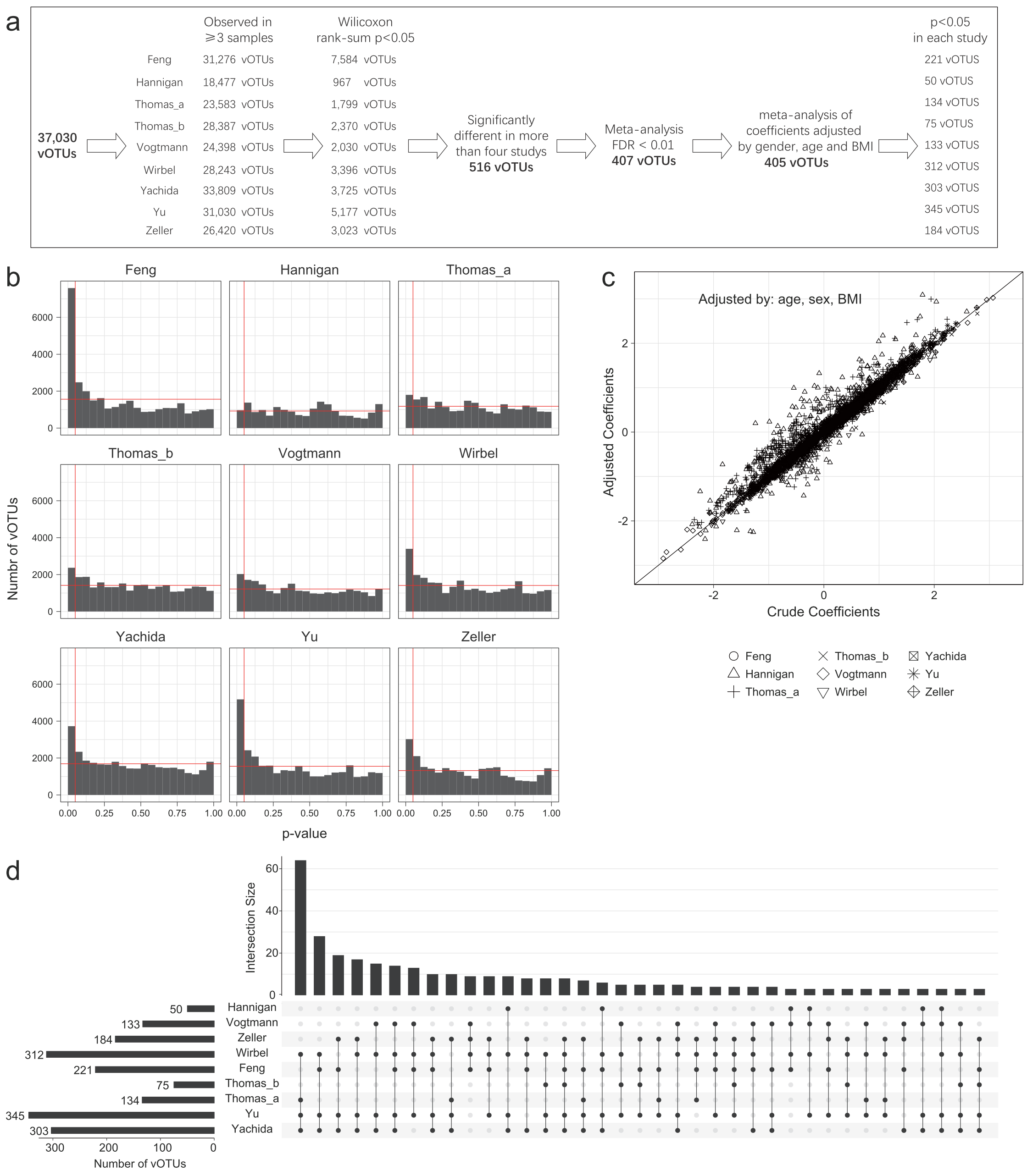

**Figure S4. CRC-associated viral biomarkers across nine studies. a**, The workflow for identifying CRC-associated viral biomarkers across nine studies. **b**, Distribution of p values from Wilcoxon rank-sum test for the relative abundance of vOTUs in each case-control study (health control versus CRC). X-axis in red, p-value at 0.05; Y-axis in red, the expected distribution of p values under the null hypothesis. **c**, Scatter plot of crude and age-, sex- and BMI-adjusted coefficients generated by linear regression analysis using relative abundances of 407 vOTUs with false discovery rate (FDR) <0.01 in random effects meta-analysis. **d**, Upset plot showing the overlap of viruses with statistical differences (p<0.05) using Wilcoxon rank-sum test on 405 CRC-associated viral profiles of each case-control dataset between different datasets.

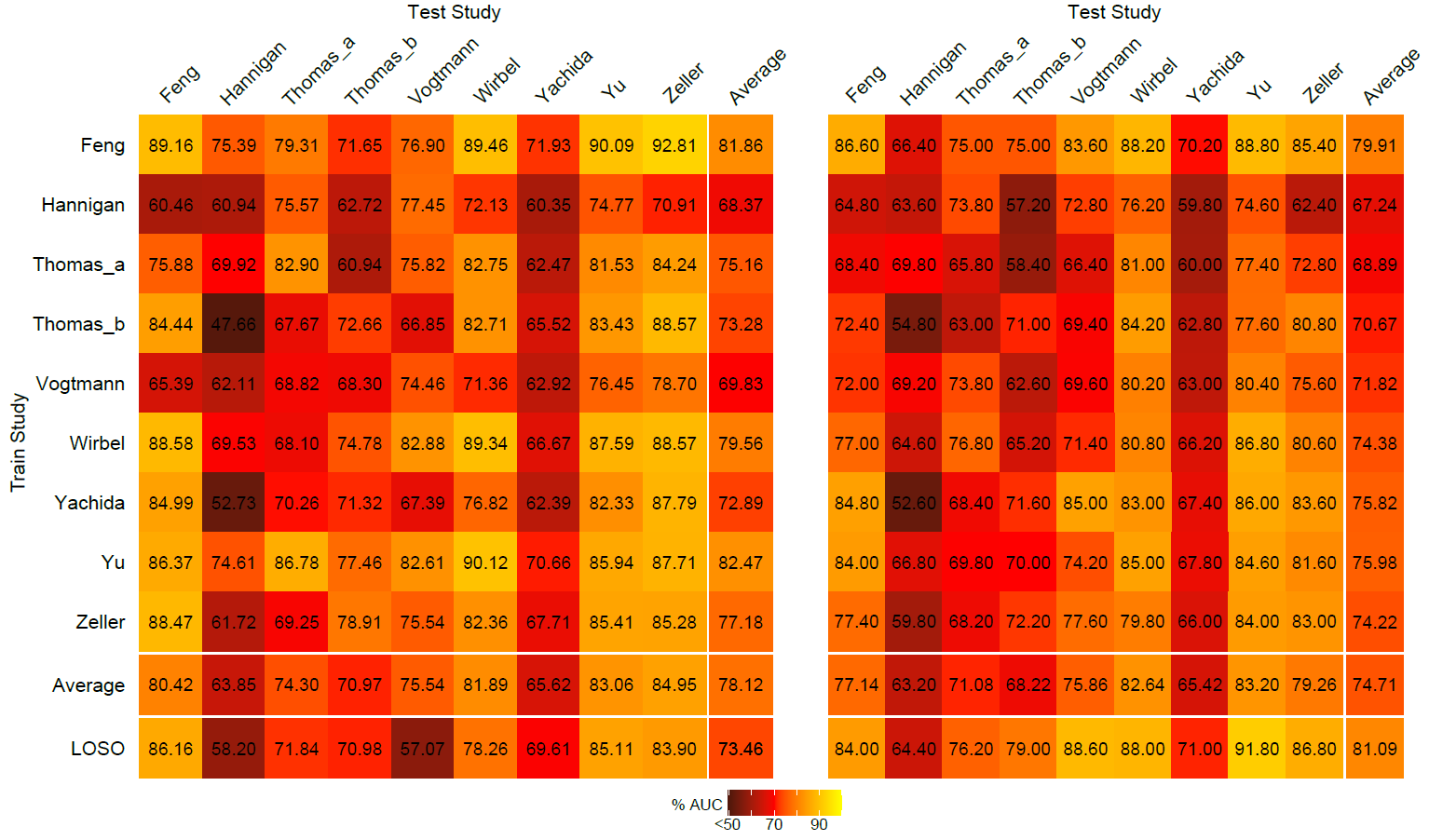

**Figure S5. Performance assessment as AUC scores of intra-cohort and cross-cohort prediction models based on relative abundances of CRC-associated vOTUs and the host properties (including age, BMI, and gender).** The model of intra-cohort prediction (diagonal) was validated using five repeats of 5-fold cross-validations. The model of cross-cohort prediction (off-diagonal) was built on the dataset corresponding to each row and validated on the dataset corresponding to each column. The LOCO row refers to leave-one-cohort-out (LOCO) analysis in which models were built on eight datasets combined and validated on the remaining one corresponding to each column. The left panel, least absolute shrinkage and selection operator (LASSO) model. The right panel, random forest (RF) model.

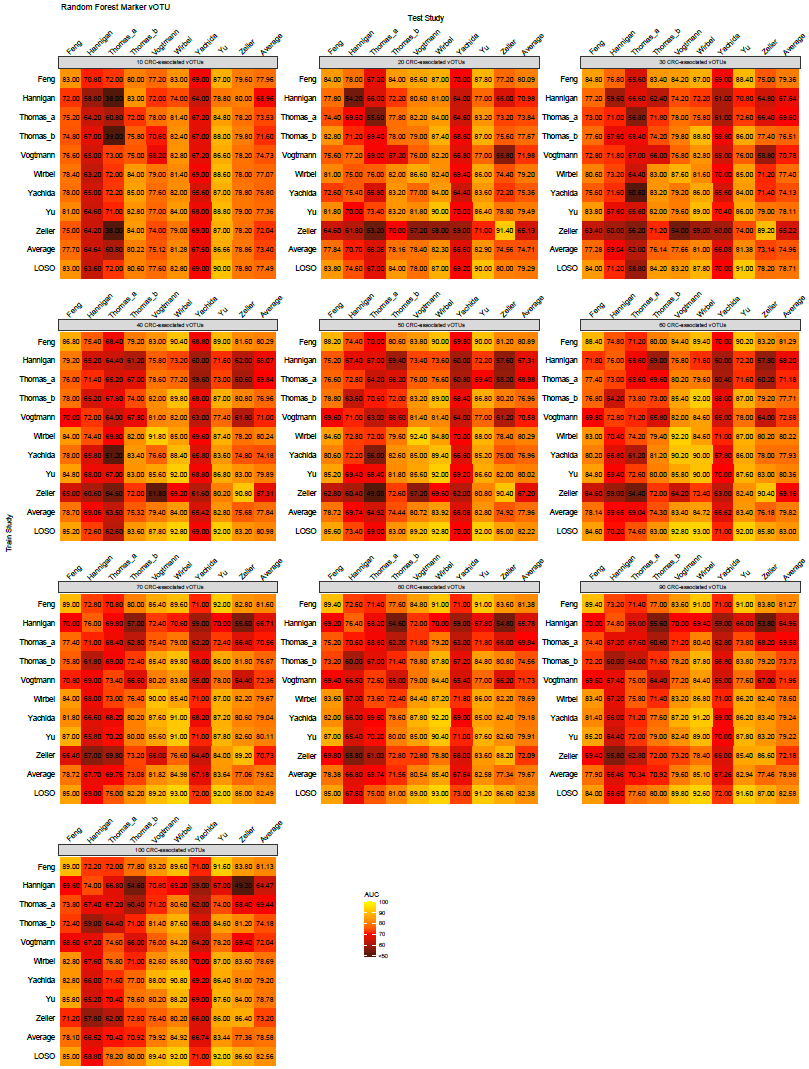

**Figure S6. Performance assessment as AUC scores for different numbers of CRC-associated vOTUs using random forest models in predicting CRC status.** The model of intra-cohort prediction (diagonal) was validated using five repeats of 5-fold cross-validations. The model of cross-cohort prediction (off-diagonal) was built on the dataset corresponding to each row and validated on the dataset corresponding to each column. The LOCO row refers to leave-one-cohort-out (LOCO) analysis in which models were built on eight datasets combined and validated on the remaining one corresponding to each column.

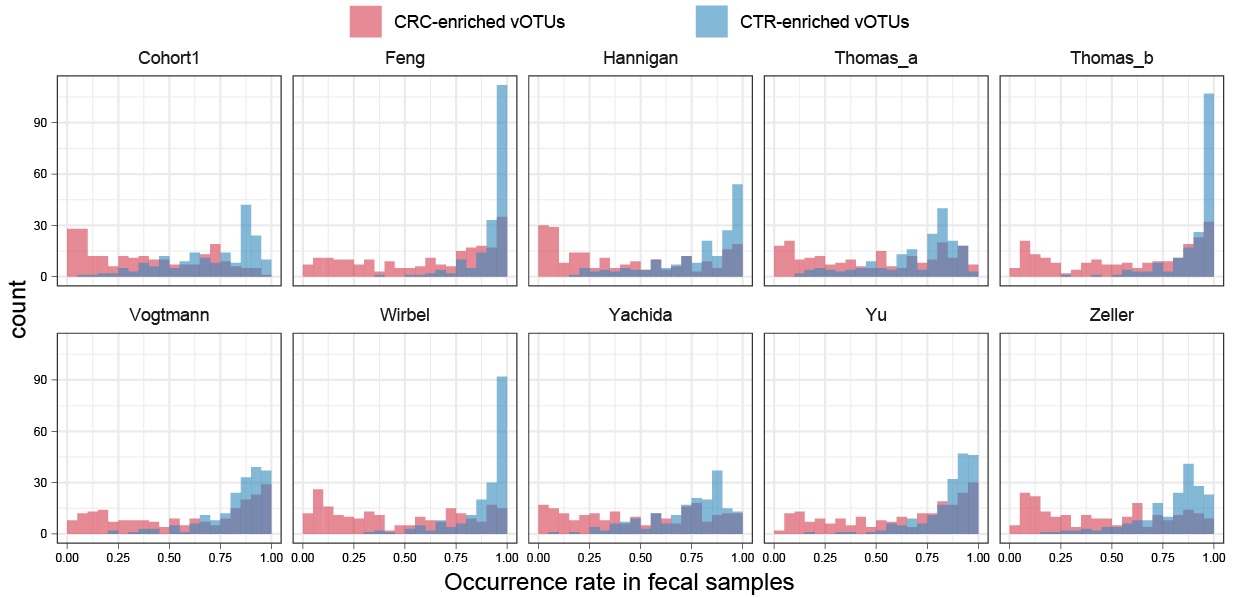

**Figure S7. Distribution of occurrence rate of CRC-associated vOTUs in fecal samples of each dataset.**
